# Modeling of axonal endoplasmic reticulum network by spastic paraplegia proteins

**DOI:** 10.1101/069005

**Authors:** Belgin Yalçın, Lu Zhao, Martin Stofanko, Niamh C O’Sullivan, Zi Han Kang, Annika Roost, Matthew R. Thomas, Sophie Zaessinger, Olivier Blard, Alex L. Patto, Valentina Baena, Mark Terasaki, Cahir J O’Kane

## Abstract

Axons contain an endoplasmic reticulum (ER) network that is largely smooth and tubular, thought to be continuous with ER throughout the neuron, and distinct in form and function from rough ER; the mechanisms that form this continuous network in axons are not well understood. Mutations affecting proteins of the reticulon or REEP families, which contain intramembrane hairpin domains that can model ER membranes, cause an axon degenerative disease, hereditary spastic paraplegia (HSP). Here, we show that these proteins are required for modeling the axonal ER network in *Drosophila*. Loss of reticulon or REEP proteins can lead to expansion of ER sheets, and to partial loss of ER from distal motor axons. Ultrastructural analysis reveals an extensive ER network in every axon of peripheral nerves, which is reduced in larvae that lack reticulon and REEP proteins, with defects including larger and fewer tubules, and occasional gaps in the ER network, consistent with loss of membrane curvature. Therefore HSP hairpin-containing proteins are required for shaping and continuity of the axonal ER network, suggesting an important role for ER modeling in axon maintenance and function.

## Introduction

Axons allow long-range bidirectional communication in neurons. They carry action potentials along their plasma membrane from dendrites and cell body to presynaptic terminals; and they transport cell components and signaling complexes using motor proteins, anterogradely and retrogradely. Another potential route for communication along axons is endoplasmic reticulum (ER); axons possess a network of ER tubules, which appears physically continuous both locally (Tsukita and Ishikawa, 1976; Villegas et al., 2014) and with ER throughout the neuron (Lindsey and Ellisman, 1985; Terasaki et al., 1994). The physical continuity of ER and its ensuing potential for long-distance communication has been likened to a “neuron within a neuron” (Berridge, 1998).

Axonal ER appears mostly tubular and smooth, with some cisternae (Tsukita and Ishikawa, 1976; Lindsey and Ellisman, 1985; Villegas et al., 2014). Some rough ER is likely to be present too: numerous mRNAs are found in both growing and mature axons (Zivraj et al, 2010; Shigeoka et al., 2016), and local axonal translation can occur in response to injury (Ben-Yaakov et al., 2012; Perry et al., 2012). However, rough ER sheets (Tsukita and Ishikawa, 1976; Villegas et al., 2014), and markers of protein export and folding that are characteristic of rough ER (Röper, 2007; O’Sullivan et al., 2012), are relatively sparse in mature axons. Axonal ER therefore likely has major roles other than protein export; these could include lipid biosynthesis (Tidhar and Futerman, 2013; Vance, 2015), calcium homeostasis and signaling (Ross, 2012), and coordination of organelle physiology (Phillips and Voeltz, 2016). The continuity of the ER network suggests that some of these roles might have long-range as well as local functions; indeed, an ER-dependent propagating calcium wave is seen after axotomy of *Caenorhabditis elegans* or mammalian dorsal root ganglion neurons (Ghosh-Roy et al., 2010; Cho et al., 2013).

A strong hint of the importance of ER in axons is found in Hereditary Spastic Paraplegia (HSP), a group of axon degeneration disorders characterized by progressive spasticity and weakness of the lower limbs (Blackstone et al., 2011; Blackstone, 2012). Mutations affecting spastin, atlastin-1, reticulon-2, REEP1 and REEP2 account for most cases of autosomal dominant “pure” HSP (Hazan et al., 1999; Zhao et al., 2001; Züchner et al., 2006; Montenegro et al., 2012; Esteves et al., 2014). These proteins share a common feature of one or two hydrophobic hairpin-loops inserted in the ER membrane, promoting ER membrane curvature in a process termed hydrophobic wedging (Voeltz et al., 2006). Proteins of the REEP and reticulon families localize preferentially to tubular or smooth ER, and their loss results in disruption of ER tubular organization (Shibata et al., 2006; Voeltz et al., 2006; Park et al., 2010; Shibata et al., 2010); they may also contribute to modeling of rough ER sheets by stabilizing their curved edges (Shibata et al., 2009).

What is the link between ER modeling and axon structure and function? HSP-causing mutations often appear to cause loss of protein expression or function (Beetz et al., 2013; Novarino et al., 2014), and the ability of hairpin-loop proteins to form homomeric and heteromeric complexes (Shibata et al., 2008), allows some point mutations to have dominant negative effects (Züchner et al., 2006; Beetz et al., 2012). Therefore loss of normal ER modeling appears to compromise axon maintenance and function. Given the roles of hairpin-loop proteins in ER modeling, we aimed to test the model that hairpin-loop-containing HSP proteins organize the axonal ER network. Since reticulon and REEP family proteins are redundantly required for most peripheral ER tubules in yeast (Voeltz et al., 2006), we focus on the requirement for these two families in axons. We previously showed that knockdown of the *Drosophila* reticulon Rtnl1 causes expansion of epidermal ER sheets, and partial loss of smooth ER marker from distal but not proximal motor axons (O’Sullivan et al., 2012). Here we show that REEP proteins have similar roles. We also show that simultaneous loss of reticulon and REEP family members leads to a range of axonal ER phenotypes, including a reduced network with fewer and larger tubules, and occasional gaps in the network. Our work implicates hairpin-loop-containing HSP proteins as important players in the axonal ER network, and suggests further models for how the network is organized.

## Results

### Two widely expressed REEP proteins, ReepA and ReepB, localize to the endoplasmic reticulum

The reticulon and REEP families of double-hairpin-containing proteins are collectively responsible for formation or maintenance of most peripheral ER tubules in yeast (Voeltz et al., 2006). We previously showed that the *Drosophila* reticulon ortholog Rtnl1 was strongly localized in axons, and that its knockdown caused partial loss of a smooth ER marker in posterior larval segmental axons (O’Sullivan et al., 2012). To test the roles of *Drosophila* REEP proteins in axonal ER localization, we first dissected the ortholog relationships between the six *Drosophila* and six human REEP proteins. Multiple sequence alignment of mammalian and *Drosophila* REEP protein sequences suggested that *CG42678* was the single *Drosophila* ortholog of mammalian *REEP1-REEP4* (Fig. 1A). *CG42678* has previously been designated *Reep1* (http://flybase.org/reports/FBgn0261564.html), but we propose the name *ReepA* to reflect its orthology to the four mammalian genes *REEP1-REEP4*. Mammalian *REEP5* and *REEP6* appeared to share two *Drosophila* orthologs, *CG8331* and *CG4960* (Fig. 1). We designated *CG8331* as *ReepB* because of its widespread expression (www.flyatlas.org; Chintapalli et al., 2007) and slower rate of evolutionary sequence divergence (Supplementary Table. 1). We excluded *CG4690* and three additional *Drosophila REEP* genes from further study, since their expression was restricted to testes and larval fat body (www.flyatlas.org; Chintapalli et al., 2007), and their faster rate of evolutionary sequence divergence (Supplementary Table. 1, and reflected in longer branch lengths in Fig. 1A), suggesting poorly conserved function.

**Fig. 1.**
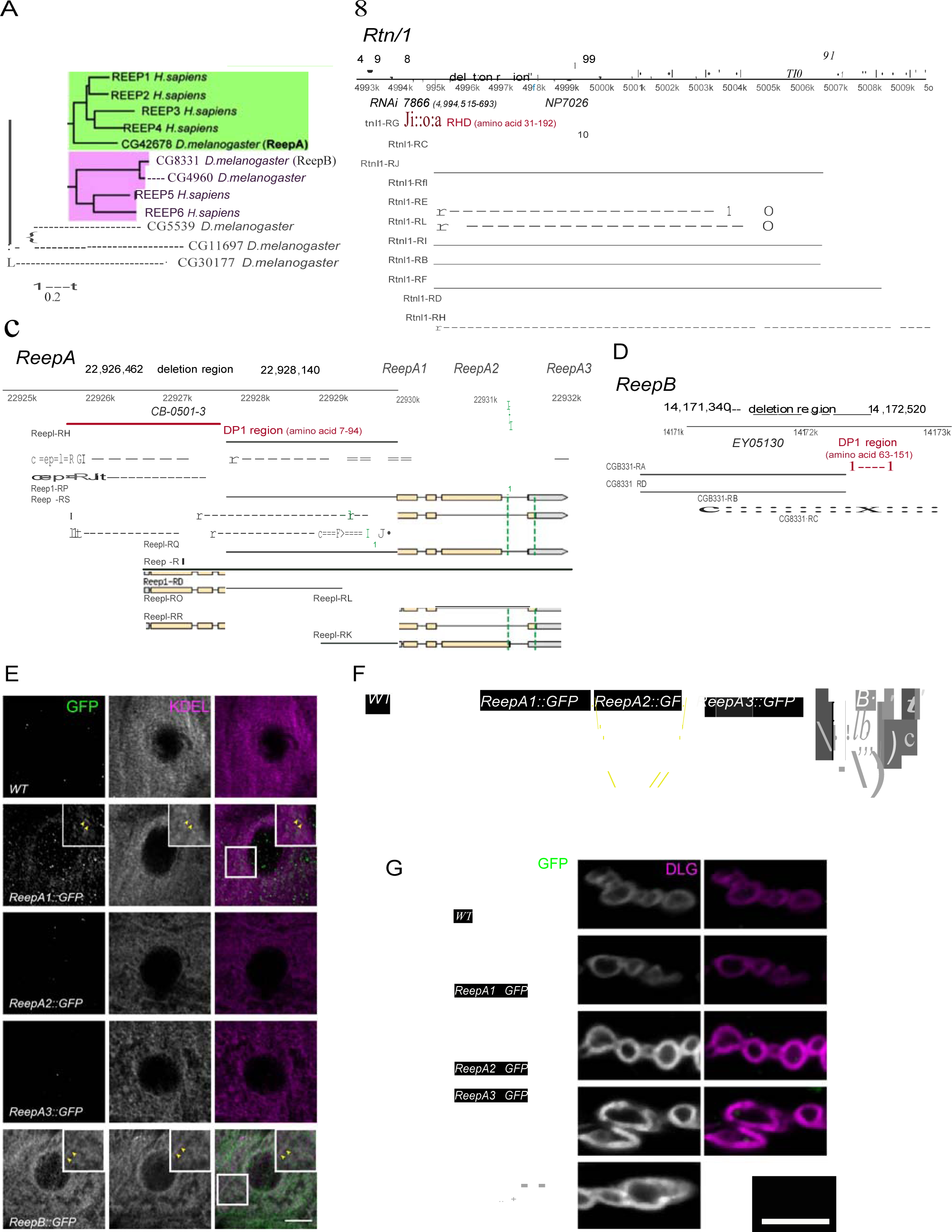
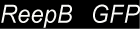
*Drosophila Rtnl1* and *REEP* genes and products. **(A)** A dendrogram based on ClustalW sequence alignment of *Drosophila* and human REEP proteins shows two branches corresponding to human REEP1-4 (CG42678) and human REEP5-6. Broken lines represent *Drosophila* REEP proteins that are evolving rapidly, reflected by longer branch lengths. Sequences used are NP_075063.1, NP_057690.2, NP_001001330.1, NP_079508.2, NP_005660.4, NP_612402.1, NP_726266.1, NP_610936.2, NP_651429.1, NP_611831.2, NP_572730.1, NP_726366.1. **(B)** *Rtnl1* genomic and transcript map, showing the region deleted in *Rtnl1*^-^ by excision of P-element *NP7026* (Wakefield and Tear, 2006; Supplementary Fig. 1), the RHD domain (Pfam 02453) and its coordinates in protein isoform G, the Rtnl1::YFP exon trap insertion *CPTI001291* (green triangle), and the fragment targeted by GD RNAi 7866. Map and coordinates are from the *Drosophila* Genome Browser (www.flybase.org, version R6.04), here and in subsequent panels; light regions in transcripts represent coding regions, dark shaded regions represent untranslated regions. **(C)** *ReepA* genomic and transcript map showing the region deleted in *ReepA*^-^ by excision of *P*-element *CB-0501-3*, the position of the DP1 domain (Pfam 03134) and its coordinates in protein isoforms H and J. GFP insertion sites for *ReepA1::GFP, ReepA2::GFP* and *ReepA3::GFP* fusions are shown with green triangles. **(D)** *ReepB* genomic and transcript map showing the region deleted in *ReepB*^-^ by excision of *P*-element *EY05130*, the position of the DP1 domain (Pfam 03134) and its coordinates in protein isoforms A and D). (**E-G**) Confocal sections showing localization of ReepA::GFP isoforms and ReepB::GFP. **(E)** Overlap of ReepA1::GFP and ReepB::GFP with anti-KDEL labeling in larval epidermal cells. To facilitate display of weaker ReepA1::GFP, the GFP channel in wild-type control (WT) and ReepA::GFP images has been brightened four times as much as for ReepB::GFP. **(F)** Expression of ReepA3::GFP and ReepB::GFP in third instar ventral nerve cord. The GFP channel for wild-type and ReepA::GFP have been brightened by twice as much as for ReepB::GFP. VNCs are outlined with the yellow dashed line. **(G)** Double labeling of ReepA::GFP and ReepB::GFP lines for GFP and Dlg (mainly postsynaptic) shows presynaptic expression of ReepB::GFP (Scale bars: E, G 10 µm; F 20 µm).

A Rtnl1::YFP exon trap, *Rtnl1*^*CPTI001291*^ (Fig. 1B) was previously shown to localize to ER, including in axons (O’Sullivan et al., 2012). To study localization of ReepA and ReepB, we recombineered C-terminal GFP-tagged versions of these (Fig. 1C,D) using P[acman] genomic clones (Venken et al., 2009). For ReepA, we generated EGFP fusions at three different C-termini (Fig. 1C), that we called ReepA1::GFP (for protein isoforms D, E, G), ReepA2::GFP (for protein isoforms H, I, J, K) and ReepA3::GFP (for protein isoforms L, R, S). In epidermal cells, we detected weak expression of ReepA1::GFP, but not of the other ReepA::GFP fusions; ReepB::GFP showed stronger expression still, and both fusions overlapped with an ER marker (KDEL; Fig. 1E), similar to REEP proteins in other organisms (Shibata et al., 2008; Park et al., 2010). ReepB::GFP was also more strongly expressed in third instar larval CNS than any ReepA3::GFP, which was the only ReepA::GFP fusion that we detected there (Fig. 1F). ReepB::GFP, but no ReepA::GFP fusion, was also detected in segmental nerve bundles emerging from the CNS (Fig. 1F), and as a mostly continual structure along the length of presynaptic terminals of neuromuscular junctions (NMJs) (Fig. 1G).

An *Rtnl1* loss-of-function mutant, *Rtnl1*^*1*^ (Wakefield and Tear, 2006), hereafter referred to as *Rtnl1*^-^, lacks the hydrophobic hairpin loop domain (RHD region) that induces ER membrane curvature (Fig. 1B; Supplementary Fig. 1A). We also used *P*-transposase-mediated imprecise excision to generate *ReepA*^-^ and *ReepB*^-^ mutants, both of which lack most of the curvature-mediating DP1 hairpin domains (Fig.1B,C; Supplementary Fig. 1B,C).

### *Rtnl1*^-^ and *ReepA*^-^ *ReepB*^-^ larval epidermal cells display an abnormal ER network

First, we asked whether *Drosophila* reticulon and REEP proteins contribute to ER network organization in third instar larval epidermal cells; *Rtnl1* knockdown leads to a more diffuse ER network organization and expansion of ER sheets in these cells, compared to wild-type controls (O’Sullivan et al, 2012). *Rtnl1*^-^ mutant larvae also showed loss of ER network organization (Fig 2A). Intensity of KDEL labeling along a line from the nucleus to cell periphery displayed fluctuating intensity, reflecting the reticular distribution of KDEL in wild-type larvae, but less fluctuation in *Rtnl1*^-^ larvae; overall levels of KDEL remained unchanged (Fig. 2A). Similarly, loss of both *ReepA* and *ReepB*, but not loss of either gene alone, made KDEL levels in larval epidermal cells fluctuate less than in controls. Mean intensity of KDEL staining was decreased in all *ReepA* and *ReepB* mutant genotypes, and fluctuation in intensity was increased in *ReepB*^-^ larvae (Fig. 2B). Ultrastructural analysis revealed that *ReepA*^-^ *ReepB*^-^ double mutant epidermal cells showed longer ribosome studded sheet ER profiles, compared to controls, and to *ReepA*^-^ and *ReepB*^-^ single mutants (Fig. 2C). This mutant phenotype is similar to, although less severe than loss of *Rtnl1* (O'Sullivan et al., 2012). *ReepA*^-^ *ReepB*^-^ mutants also showed increased ER stress in epidermal cells (Fig. 2D) but not in CNS (Fig. 2E). Therefore, Rtnl1 and REEP proteins shape the ER network in *Drosophila*, and their loss disrupts ER organization, causing longer ER sheet profiles.

**Fig. 2.**
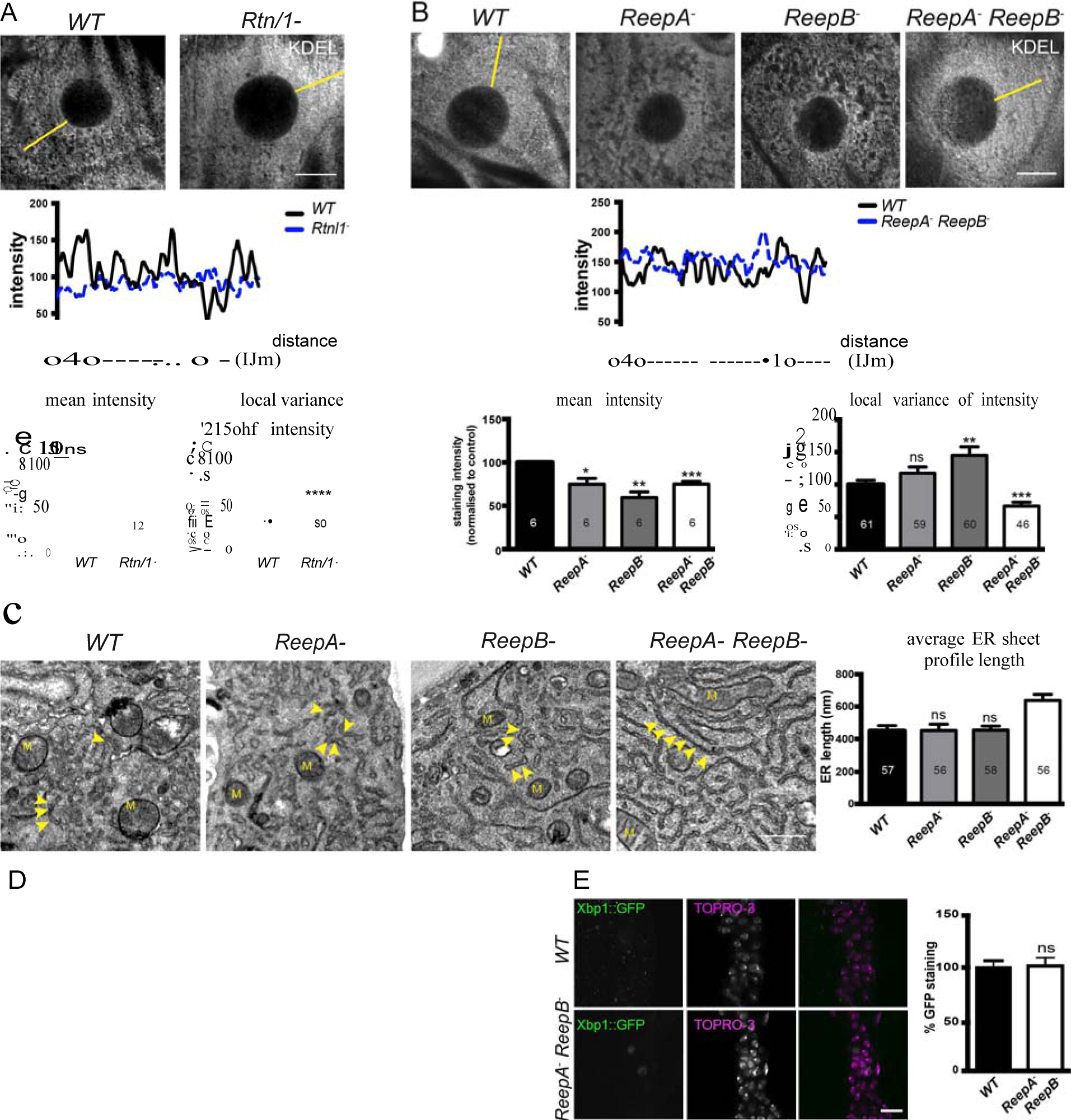
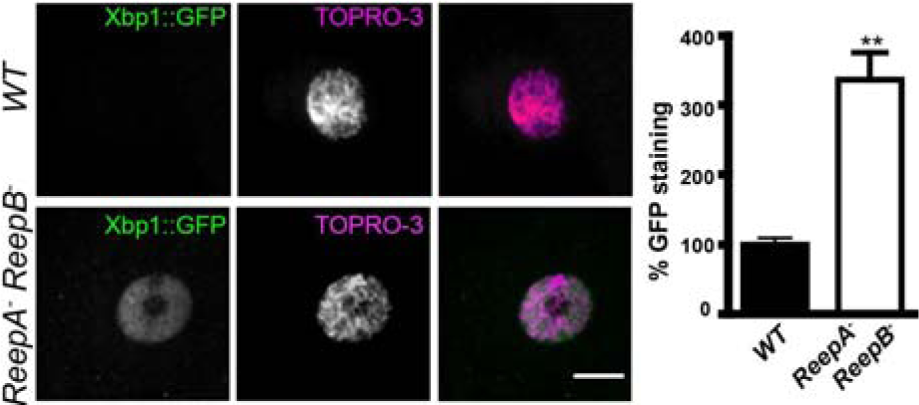
*Rtnl1*^-^ mutants and *ReepA*^-^ *ReepB*^-^ double mutants show disrupted ER organization in epidermal cells. **(A)** KDEL distribution in third instar larval epidermal cells appears diffuse in *Rtnl1*^-^ larvae compared to a more reticular staining in wild-type (WT) larvae. Micrographs show 1.5-µm z-projections of confocal sections. KDEL intensity along yellow lines from nuclear envelope to the cell periphery shows less fluctuation in *Rtnl1*^-^ (blue dotted line on line graph) than in *WT* larvae (black line on line graph). Fluctuation of intensity is quantified using the local normalized variance of intensity (variance/mean for rolling 10-pixel windows) over a 12-µm line for each cell. n=61 epidermal cells from 12 different larvae, 3-5 cells from each larva, from 3 independent experiments. **(B)** KDEL distribution in third instar larval epidermal cells shows less spatial fluctuation in *ReepA*^-^ *ReepB*^-^ double mutant larvae compared to wild-type (*ReepA*^+^), and *ReepA*^-^ and *ReepB*^-^ single mutant larvae. This is confirmed by quantification as in **A**; overall KDEL intensity is reduced in all ReepA and ReepB mutant genotypes. n=45-61 epidermal cells in total from 12-16 different larvae, 3-5 cells from each larva, from 6 independent experiments. **(C)** Electron micrographs show increased ER sheet profile length in *ReepA*^-^ *ReepB*^-^ double mutant third instar larval epidermal cells compared to single *ReepA*^-^ and *ReepB*^-^ mutants and controls. Arrowheads, ER sheets; M, mitochondria. (n=56-58 cells in total from 3 independent larvae, 18-19 cells from each. **(D,E)** *ReepA*^-^ *ReepB*^-^ double mutant larvae have an increased ER stress response compared to a *ReepA*^+^ wild-type control, measured by Xbp1::GFP expression, in larval epidermis **(D)** but not in neuronal cell bodies **(E)**. Graphs show quantification of Xbp1::GFP staining intensity relative to controls. (n = 12-15 larvae). All bar graphs show mean ± SEM. ns, P > 0.05; *, P < 0.02; **, P < 0.005; ***, P < 0.0003; ****, P < 0.0001, two-tailed Student’s t-test. Scale bars: A, B 10 µm; C, 0.5 µm).

### Loss of Rtnl1 or ReepB causes partial loss of axonal ER marker from posterior axons

To understand the roles of reticulon and REEPs in axonal ER organization, we labeled axonal ER by expressing Acsl::myc (O’Sullivan et al., 2012) in two adjacent motor neurons using *m12-GAL4* (Xiong et al. 2010). Loss of *Rtnl1* caused partial loss of Acsl::myc from posterior (segment A6) axons, but not from anterior axons (segment A2); it also caused Acsl::myc staining to appear more irregular in posterior but not anterior axons, reflected in a higher coefficient of variation (SD/mean) of Acsl::myc staining intensity along the length of posterior axons lacking Rtnl1, compared to wild-type (Fig. 3A). These *Rtnl1* loss-of-function phenotypes were found either on targeted knockdown of Rtnl1 in these motor axons or in *Rtnl1*^-^ mutant larvae, and the *Rtnl1*^-^ mutant phenotypes could be partially rescued by one copy of an *Rtnl1*^*Pacman*^ genomic clone (Fig. 3A).

**Fig. 3.**
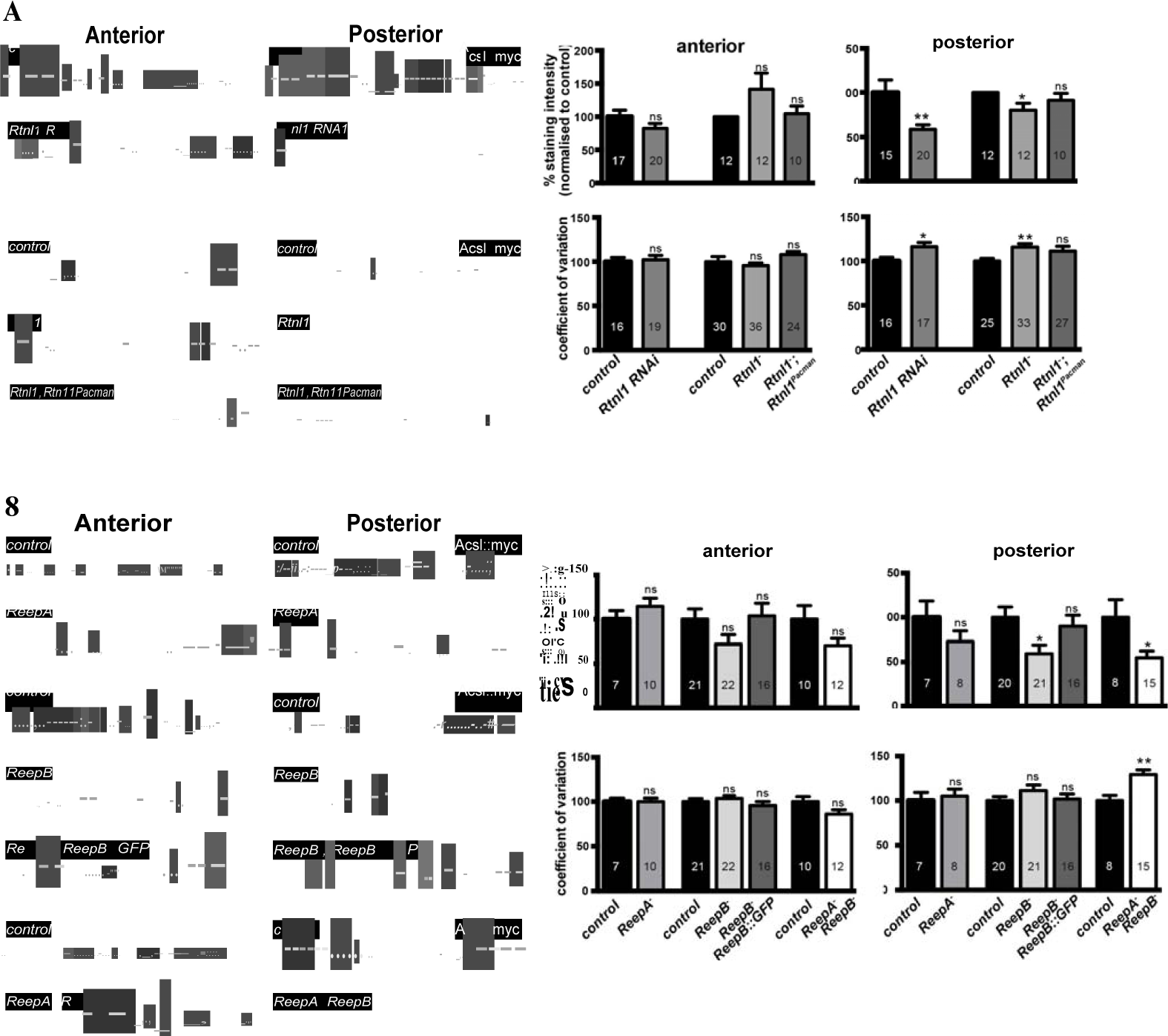
Loss of either Rtnl1 or ReepB leads to partial loss of smooth ER marker from distal motor axons. **(A)** Effects of Rtnl1 loss by RNAi knockdown (top images) or *Rtnl1*^-^ mutant (bottom images) on ER visualized by Acsl::myc expressed in two adjacent motor axons using *m12-GAL4*. Rtnl1 loss leads to partial loss of Acsl::myc in posterior but not anterior axons (top graphs), and to some disorganization of posterior axonal ER seen by increased coefficient of variation of Acsl::myc staining intensity. There is partial rescue of *Rtnl1* phenotypes by one copy of a *Rtnl1*^*Pacman*^ genomic clone. n=15-20 larvae per genotype pooled from 5 independent experiments for RNAi; n=24-36 larvae per genotype from 10-12 independent experiments for *Rtnl1*^-^ mutant. **(B)** Loss of *ReepB* but not of *ReepA* causes partial loss of smooth ER marker Acsl::myc in posterior but not in anterior motor axons. This phenotype can be partially rescued by one copy of a genomic *ReepB::GFP* clone. A *ReepA*^-^ *ReepB*^-^ double mutant shows a similar phenotype to a *ReepB*^-^ single mutant. *ReepA*^-^ *ReepB*^-^ double mutant, but not single *ReepA*^-^ or *ReepB*^-^ mutants, show increased coefficient of variation of Acsl::myc staining levels in posterior axons; n=7-22 larvae from 4-11 independent experiments. All graphs show mean ± SEM; ns, P > 0.05; *, P < 0.03; **, P < 0.005 except where otherwise indicated; two-tailed Student’s t-test. Scale bars, 10 µm.

*ReepA*^-^ mutants showed no loss of Acsl::myc from axons. Similar to *Rtnl1* loss of function, *ReepB*^-^ mutants showed partial loss of Acsl::myc from posterior but not anterior motor axons (Fig. 3B); this phenotype was partially rescued with one copy of a genomic *ReepB::GFP* clone. A *ReepA*^-^ *ReepB*^-^ double mutant showed loss of Acsl::myc from posterior axons, that was similar to that seen in *ReepB*^-^ mutants (Fig. 3B). *ReepA*^-^ *ReepB*^-^ double mutant larvae, but not *ReepA*^-^ or *ReepB*^-^ single mutants, showed an increased coefficient of variation of Acsl::myc staining intensity in posterior but not in anterior axons (Fig. 3B). In summary loss of either Rtnl1 or at least ReepB alters axonal ER distribution in posterior motor axons, therefore supporting roles for these two protein families in axonal ER organization.

### *Rtnl1*^-^ and *Rtnl1*^-^ *ReepA*^-^ *ReepB*^-^ mutants disrupt axon ER integrity and axon transport

Since loss of both reticulon and REEP families in yeast removes most peripheral ER tubules (Voeltz et al., 2006), we tested whether loss of both protein families in *Drosophila* might have similarly severe effects on axonal ER. *Rtnl1*^-^ *ReepA*^-^ *ReepB*^-^ triple mutant larvae showed increased fluctuation of Acsl::myc staining intensity along motor axons compared to wild-type, mainly in middle parts (segment A4 and A5) of longer axons. At its most extreme, this manifested as fragmentation of Acsl::myc labeling, which was never seen in wild-type axons (Fig. 4A). When gaps in labeling were found in the central regions of axons, labeling in the anterior and posterior parts of the same axons was usually continuous. Double labeling of plasma membrane (CD4::tdGFP) and axonal ER (Acsl::myc) showed no effect on axonal plasma membrane in *Rtnl1*^-^ *ReepA*^-^ *ReepB*^-^ triple mutant compared to control axons (Fig. 4B), implying that the phenotype was limited to ER and did not affect axon integrity. To compare genotypes and labels, we quantified irregular labeling along a 45-µm stretch of axon traversing the larval A4/A5 region in two ways: first we measured gaps in labeling, defined as intensity below a threshold that could consistently distinguish labeled axons above background labeling in the nerve; and second, we quantified the coefficient of variation of labeling intensity along the axon (Fig 4B,C). *Rtnl1(RNAi) ReepA*^-^ *ReepB*^-^ larvae also showed a fragmentation phenotype similar to *Rtnl1-ReepA-ReepB-*, suggesting that loss of Rtnl1 is essential for it (Fig. 4C). *Rtnl1* loss-of-function axons, but not *ReepA*^-^ *ReepB*^-^ mutant axons, also showed a mild ER fragmentation phenotype (Fig 4A,C). Therefore, loss of Rtnl1 causes a mild irregular organization of axonal ER, and this phenotype is exacerbated by loss of Reep proteins; however, loss of Reep proteins alone has no apparent effect.

**Fig. 4.**
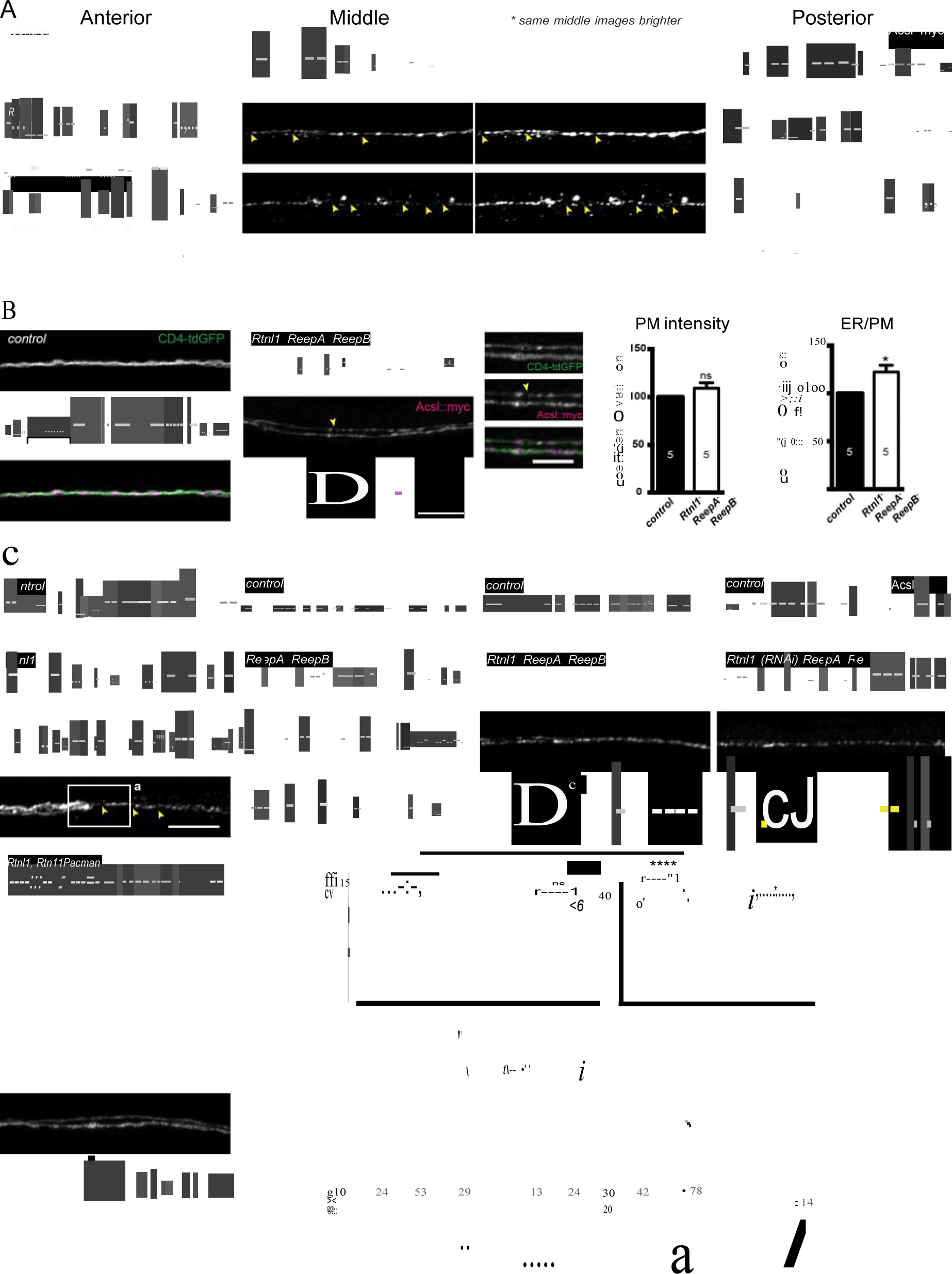
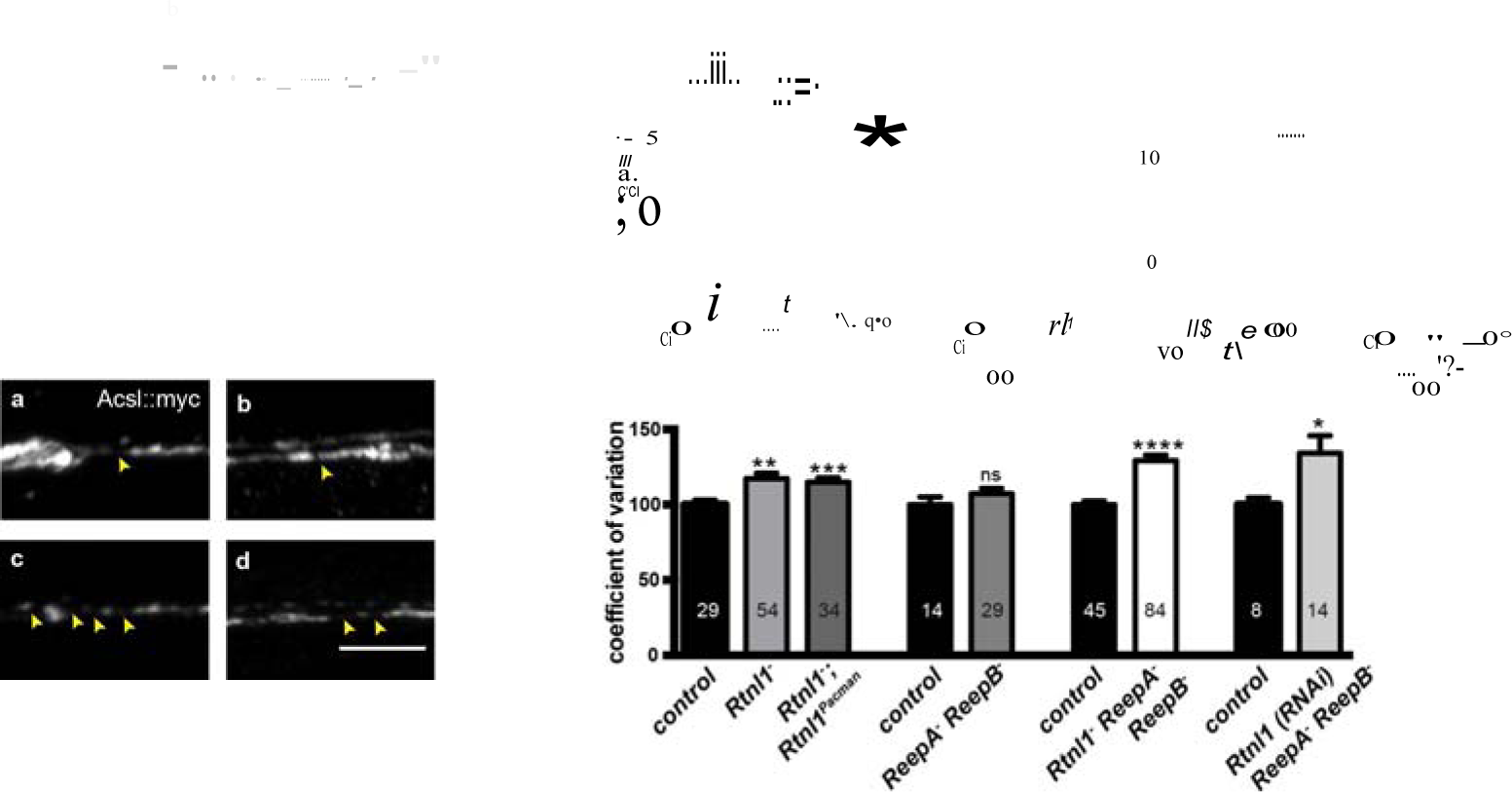
Loss of hairpin proteins leads to discontinuity of axonal ER staining. **(A)***Rtnl1*^-^ larvae and *Rtnl1*^-^ *ReepA*^-^ *ReepB*^-^ triple mutant larvae sometimes show fragmented axonal ER labeling in the middle parts (segment A5) of long motor axons that express Acsl::myc under control of *m12-GAL4*. Anterior (segment A2) and posterior (segment A6) portions of the same motor axons show continuous ER labeling. Arrowheads show gaps in Acsl::myc staining; brighter versions of the same images show gaps in staining in mutants but not wild-type, even when brightness of remaining staining is saturating. **(B)** Coefficient of variation for plasma membrane labeling along axon length is unchanged in *Rtnl1*^-^ *ReepA*^-^ *ReepB*^-^ triple mutant compared to control axons, whereas the ER to plasma membrane ratio of the coefficient of variation is increased. Higher zoom images of the boxed areas show gaps (yellow arrowheads) and brightly labeled axonal ER accumulations. Graphs show mean ± SEM; *, P<0.02, two-tailed Student’s t-test; n=6 different larvae from 3 independent experiments. **(C)** A variety of phenotypes, from continuous to severely fragmented Acsl::myc labeling with long gaps, are found in *Rtnl1*^-^ larvae, *Rtnl1*^-^ *ReepA*^-^ *ReepB*^-^ triple mutant larvae, and *Rtnl1(RNAi) ReepA*^-^ *ReepB*^-^ larvae. Top graphs show percentages of a 45-µm length in the middle (A4/A5 segment) of each axon that lacks Acsl::myc staining, using an intensity threshold of 20 on a scale of 0-255. Individual axons are plotted, together with median and interquartile range; comparisons use Mann-Whitney U-tests; two datapoints are above the top of the scale in the *Rtnl1*^-^ data. The coefficient of variation of Acsl::myc labeling in middle axon portions is also increased in these genotypes relative to controls (bottom graph, mean ± SEM, Student’s t test). In all experiments, except where stated: ns, P > 0.05; *, P < 0.05; **, P < 0.005; ***; P < 0.001; ****, P < 0.0001; n=3-11 independent experiments, each, with 2-3 different larvae for each genotype. Scale bars 10 µm, and 5 µm in higher zoom images).

We also tested for possible defects in axon transport by staining for abnormal accumulation of the synaptic vesicle protein CSP in axons. *Rtnl1*^-^ larvae and *Rtnl1*^-^ *ReepA*^-^ *ReepB*^-^ triple mutant larvae showed large accumulations of CSP in many peripheral nerves. The large accumulations of CSP in *Rtnl1*^-^ larvae could be rescued by two copies of a *Rtnl1*^*Pacman*^ genomic clone (Fig. 5).

**Fig. 5.**
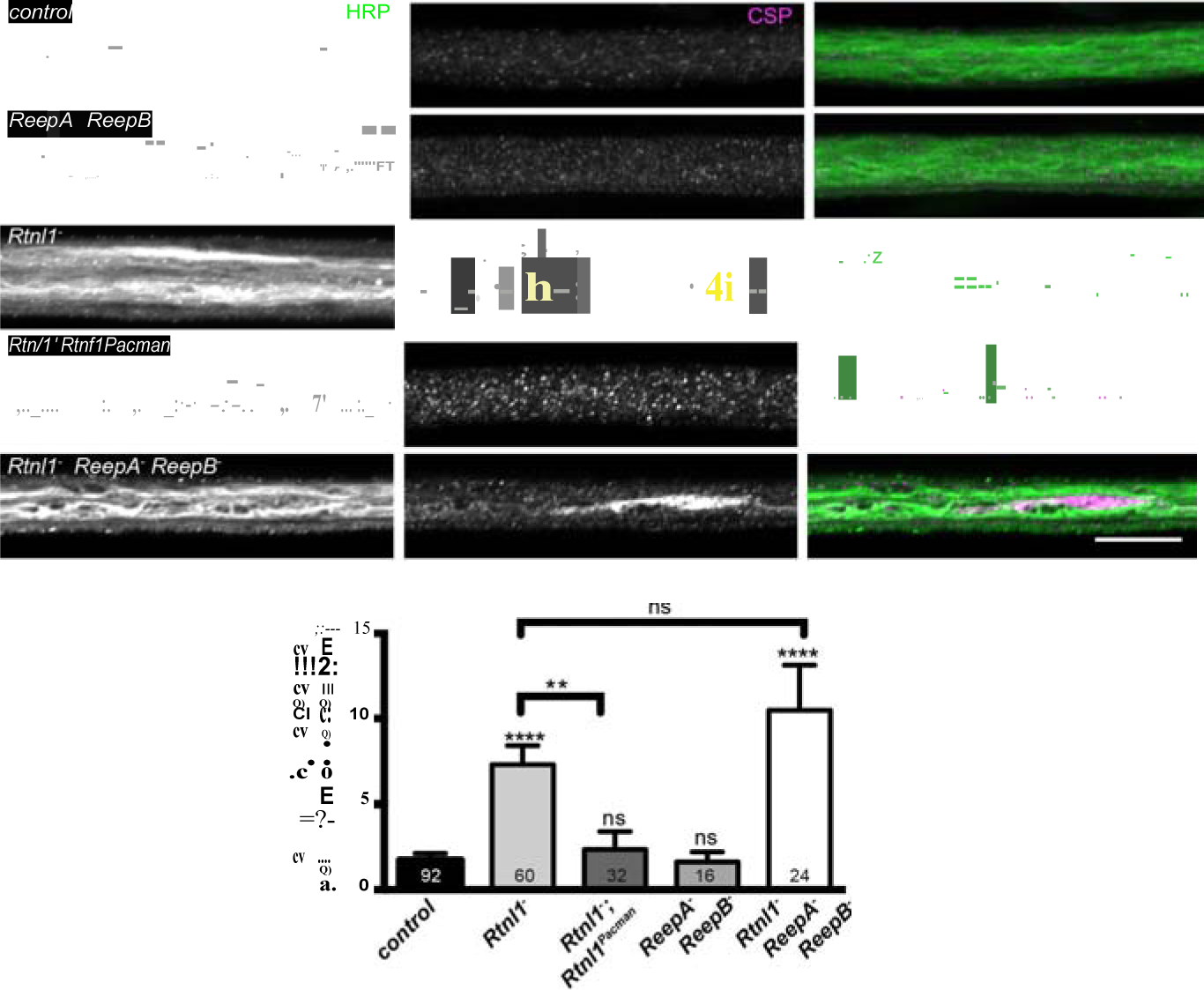
Loss of Rtnl1 causes mild accumulation of synaptic vesicles in axons. Peripheral nerves of *Rtnl1*^-^ and *Rtnl1*^-^ *ReepA*^-^ *ReepB*^-^ triple mutant larvae show larger accumulations of synaptic vesicle protein CSP (e.g. yellow arrows), and smaller elongated CSP puncta (e.g. yellow arrowheads). In contrast, control larvae and *ReepA*^-^ *ReepB*^-^ double mutant larvae show an even distribution of small round CSP puncta. CSP accumulations are not significantly bigger in *Rtnl1*^-^ *ReepA*^-^ *ReepB*^-^ triple mutants than in *Rtnl1*^-^ mutants. The CSP accumulations in *Rtnl1*^-^ larvae can be rescued by two copies of a *Rtnl1*^*Pacman*^ genomic clone. All axons shown are crossing abdominal segment A2. Graph shows mean ± SEM; n = 16-92 axons from 8-46 larvae, from 3 different experiments. ns, P > 0.05; **, P < 0.006; ****, P < 0.0001, two-tailed Student’s t-test. Scale bar, 10 µm).

### *Rtnl1*^-^ *ReepA*^-^ *ReepB*^-^ mutant nerves show ER abnormalities in axons and glia

To better understand wild-type axonal ER organization, and the mutant phenotypes seen in confocal microscopy, we performed electron microscopy (EM) on 60-nm-thick serial sections of third instar peripheral nerves. Peripheral nerves contain both motor and sensory axons, arranged in fascicles, and wrapped in three main classes of glial cell (Stork et al., 2008; Matzat et al., 2015). We used ROTO staining (Tapia et al., 2012; Terasaki et al., 2013) to preferentially highlight cellular membranes including ER.

Wild type larvae showed a network of ER tubules, in every axon that could be observed (Fig. 6A left; serial sections in Supplementary Movie 1). Tubule outer diameter averaged around 40 nm (Fig. 6B,C), and axons contained an average of around 1.6 ER tubules in each cross-section (Fig. 6D,E). Reconstruction (Fig. 6F left; Supplementary Movie 2) showed a tubular network with multiple branches, some dead ends, and continuity along nearly every axon sectioned. ER tubules often showed proximity to mitochondria or plasma membrane (Fig. 6G left), We also found occasional structures resembling small patches of ER sheets with an adjoining cisterna (Fig. 6H), continuous with the tubular ER network (Supplementary Movie 3), at a frequency averaging around one per 20 µm per axon.

**Fig. 6.**
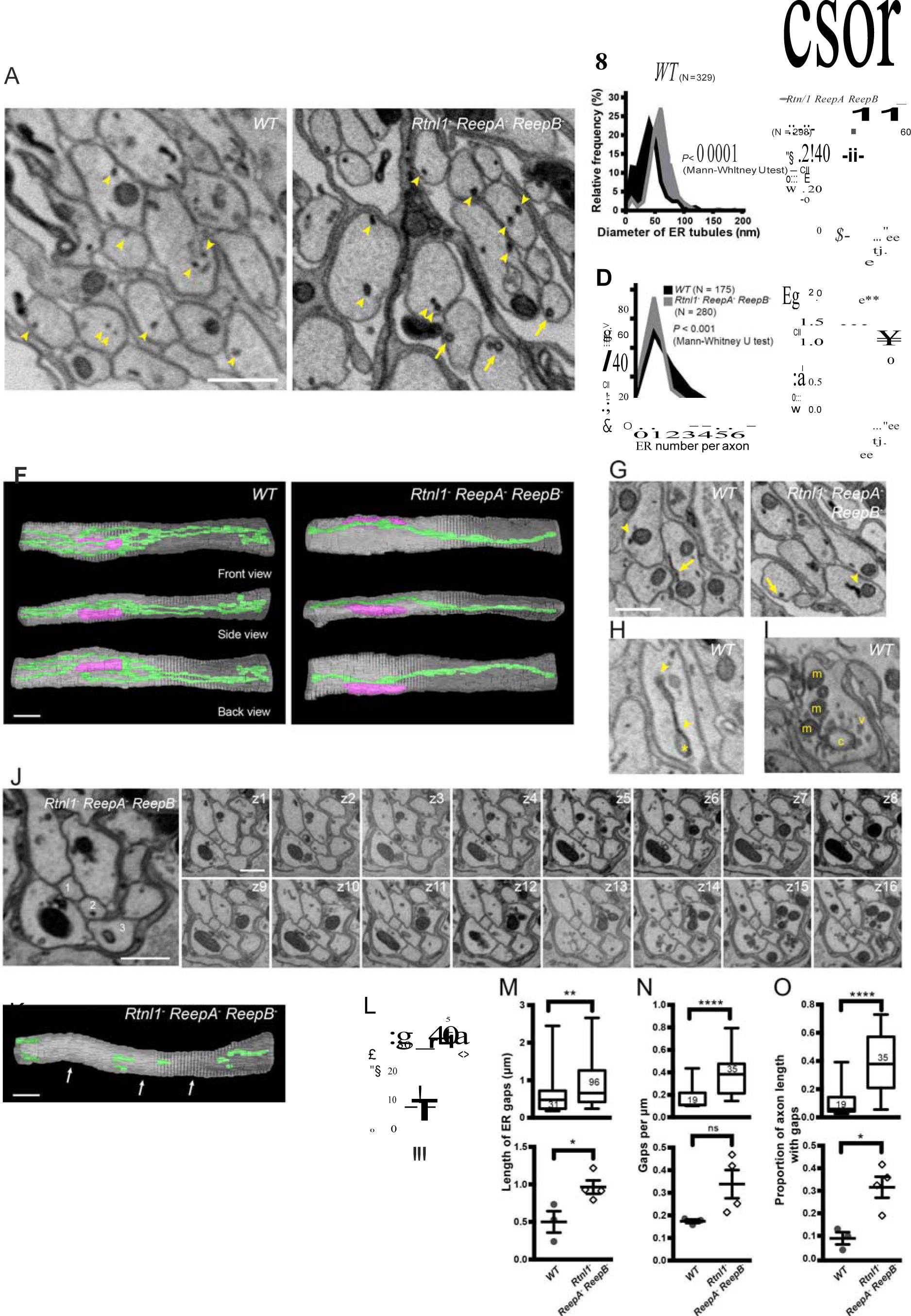
Loss of hairpin proteins leads to fewer but enlarged ER tubules in larval peripheral nerve axons. **(A)**EMs of peripheral nerve axons from wild-type (*ReepA*^+^, left) or *Rtnl1*^-^ *ReepA*^-^ *ReepB*^-^ triple mutant (right) larvae. Arrowheads indicate ER tubules (seen as continuous structures in serial sections in Supplementary Movies 1 and 4). Triple mutant axons show enlarged ER tubules, sometimes with a clear lumen (arrows), seldom observed in wild-type. Quantification of ER tubule diameter (**B,C**) and tubules per axon cross-section (**D,E**) in control and mutant larvae. Data from individual ER tubules or axons are shown in **B** and **D**, averaged larval values, mean ± SEM in **C** and **E**. **(F)** 3D reconstruction of a 4.5-µm axon segment from wild-type (left) or mutant (right) peripheral nerves, generated from 75 serial 60-nm sections, showing ER (green), mitochondria (magenta) and plasma membrane (gray). Interactive versions of the reconstructions are in Supplementary Movies 2 and 5. Some ER cisternae (arrows) are seen in the mutant axon. **(G)** Electron micrographs of peripheral nerve axons from wild-type (*ReepA*^+^, left) or *Rtnl1*^-^ *ReepA*^-^ *ReepB*^-^ triple mutant (right) larvae, showing proximity of ER to mitochondria (arrowheads) or plasma membrane (arrows). **(H)** Representative EM of short ER sheet (arrowhead) and cisterna (asterisk) from a wild-type larva; further sections in Supplementary Movie 3. **(I)** section of an axonal swelling from a wild-type larva showing mitochondria (m), vesicles (v), and a large clear cisterna (c); further sections in Supplementary Movie 6. **(J)** Serial EM sections show ER discontinuity in two mutant axons: axon 1 lacks ER tubules in sections z6-z14 and axon 3 in sections z1-z11; neighboring axons (e.g. axon 2) show a continuous ER network. **(K)** 3D reconstruction of a 4.5-µm axon segment from mutant peripheral nerves, generated from 75 serial 60-nm sections, showing multiple gaps (indicated by arrows). ER is in green and plasma membrane in gray. Raw EM data and an interactive version of the reconstruction are in Supplementary Movies 7 and 8, respectively. **(L)** Frequency of axons with gaps in the 4.5-µm lengths analyzed. ER gap length **(M)**, numbers of ER gaps per µm **(N)**, and proportion of axon length with gaps **(O)** in affected axons. In **M**-**O**, top graphs show data from individual axons, with second and third quartiles and 5th and 95th percentiles; bottom graphs show averaged larval values, mean ± SEM. In all graphs, ns P > 0.05; *, P < 0.04; **, P < 0.003; ****, P < 0.0001; Mann-Whitney U test for **B**, **D**, top graphs in **M**-**O**; two-tailed Student’s T test for **C**, **E**, **L**, bottom graphs in **M**-**O**. Scale bars, 500 nm.

If *Rtnl1*^-^ *ReepA*^-^ *ReepB*^-^ triple mutant larvae have less ER membrane curvature, we would expect them to have larger ER tubules, fewer tubules per section, loss of tubules, or a combination of these. EM revealed all these mutant phenotypes to varying degrees (Fig. 6). ER tubule diameter was increased to around 60 nm (Fig. 6B,C), allowing a lumen to be seen in some tubules (Fig. 6A right; Supplementary Movie 4) that was rarely seen in wild-type (Fig. 6A left; Supplementary Movie 1), and most triple mutant axons exhibited only a single ER tubule (Fig. 6D,E). Reconstructions (Fig. 6F right; Supplementary Movie 5) showed a less extensive ER network in mutant axons. Frequent contacts of ER with mitochondria and plasma membrane were found in both wild-type and *Rtnl1*^-^ *ReepA*^-^ *ReepB*^-^ mutant axons (Fig. 6G). Both wild-type and mutant axons showed swellings containing mitochondria and clusters of vesicles resembling synaptic vesicles (Fig. 6I; Supplementary Movie 6).

Serial EM sections also revealed variable fragmentation of ER in *Rtnl1*^-^ *ReepA*^-^ *ReepB*^-^ mutant axons (Fig. 6J,K), consistent with that seen using confocal microscopy (Fig. 4). Some discontinuity of the ER network was observed in about 10% of wild-type or *Rtnl1*^-^ *ReepA*^-^ *ReepB*^-^ mutant axons (Fig. 6L). However, gaps in *Rtnl1*^-^ *ReepA*^-^ *ReepB*^-^ mutant axons (Fig. 6J, K; Supplementary movies 7,8) were longer (Fig. 6M), and slightly more numerous (but not significantly when averaged across larvae; Fig. 6N) than in wild-type axons, resulting in nearly a four-fold increase in the length of affected axons that lacked ER tubules (Fig. 6O).

*Rtnl1*^-^ *ReepA*^-^ *ReepB*^-^ mutant peripheral nerves also showed glial cell phenotypes. Wild-type peripheral nerves are surrounded by an outer perineurial glial cell, and just beneath this a subperineurial glial cell; axons or axon fascicles are wrapped imperfectly by a wrapping glia cell (Stork et al., 2008; Matzat et al., 2015). All glial classes, but particularly subperineurial glia, showed a trend towards increased ER sheet profile length compared to control cells (Fig. 7A-I), similar to *Rtnl1* knockdown (O’Sullivan et al., 2012) or *ReepA*^-^ *ReepB*^-^ double mutant (Fig. 2) epidermal cells. Triple mutant wrapping glia also displayed more extensive wrapping, sometimes completely ensheathing axons, which was rarely observed in control nerve sections (Fig. 7J-M; Supplementary Fig. 2).

**Fig. 7.**
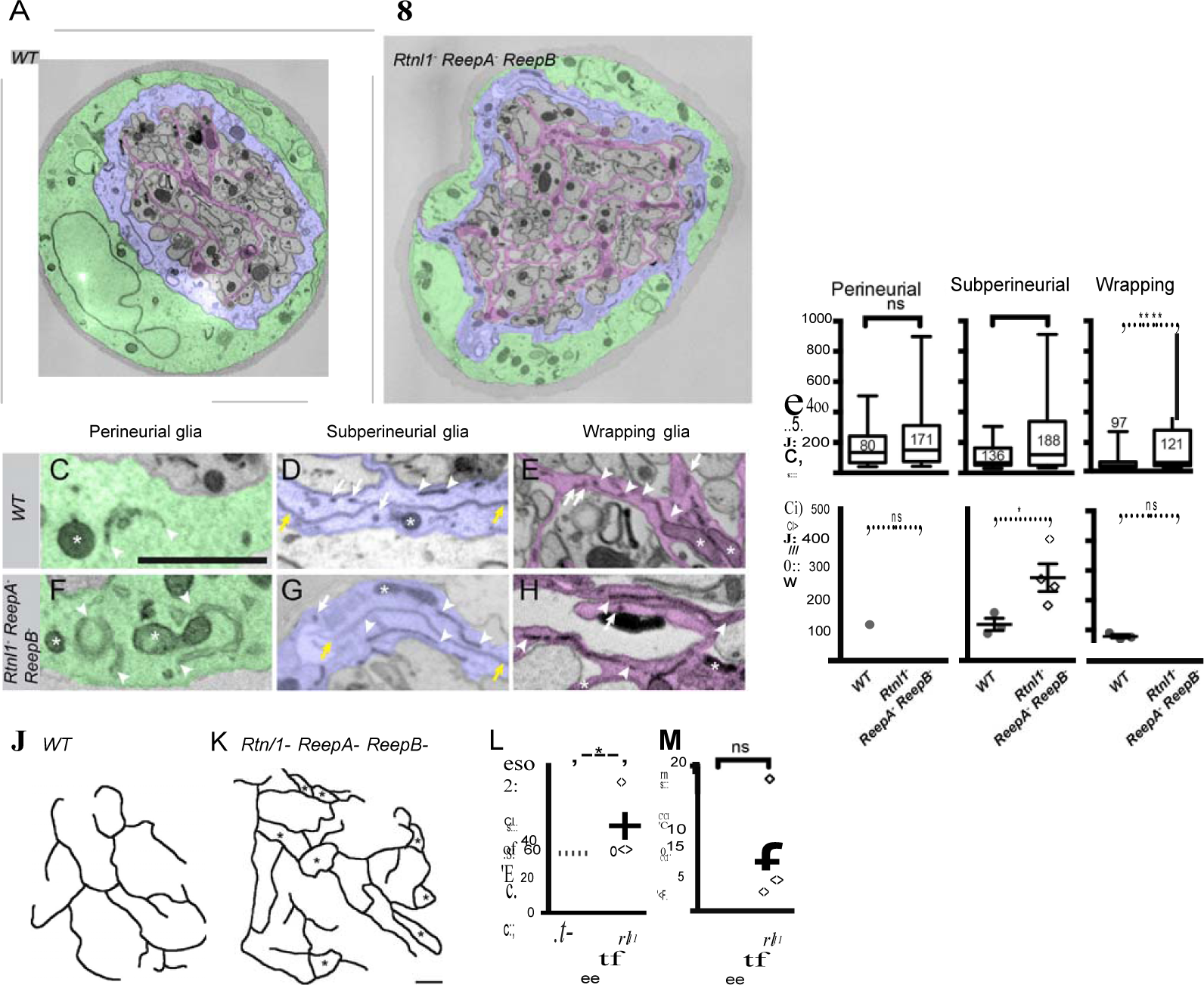
Loss of reticulon and REEP proteins leads to ER disorganization in glial cells and hyper-wrapping of peripheral axons. (**A-B**) EMs of peripheral nerve sections from wild-type (*ReepA*^+^) (**A**) or *Rtnl1*^-^ *ReepA*^-^ *ReepB*^-^ triple mutant (**B**) larvae. Perineurial, subperineurial, and wrapping glial cells are shaded green, blue, and magenta, respectively. (**C-H**) Higher magnification images of perineurial (**C,F**), subperineurial (**D,G**), and wrapping (**E,H**) glia from wild-type (**C-E**) or mutant (**F-H**) nerve sections, showing ER tubules (white arrows; confirmed as tubules by presence in adjacent sections) and sheets (white arrowheads). Note the longer ER sheet profiles and fewer ER tubules in the subperineurial (**G**) and wrapping glial cells (**H**) of mutant nerves. Asterisks show mitochondria; yellow arrowheads show glial plasma membrane, identified by its continuity. (**I**) ER sheet profile length in control and mutant glial cells from 3 wild-type and 4 mutant larvae. Top graphs represent all individual sheet profiles, with second and third quartiles and 5th and 95th percentiles; bottom graphs show averaged larval values, showing mean ± SEM. Two-way ANOVA showed a significant effect of genotype (P < 0.002) but not glial class (P > 0.3) on ER sheet length, with no interaction between factors (P > 0.3). (**J,K**) Sketches of wild-type (**J**) and mutant (**K**) wrapping glial cells, showing excess processes in the triple mutant. Asterisks indicate completely wrapped one-axon or two-axon fascicles, rarely seen in wild-type nerve sections. More examples of each phenotype are in Supplementary Fig. 2. (**L-M**) Quantification of wrapping glial membrane profile length per nerve cross-section (**L**) and percentage of axons that are wrapped individually or as two-axon fascicles (**M**) in wild-type and mutant nerve sections (mean ± SEM). ns, P > 0.05; *, P < 0.05; ***, P < 0.001; ****, P < 0.0001. Mann-Whitney U test (**I**, top graphs); two-tailed Student’s t-test (**I**, bottom graphs; **L,M**,). Scale bars, 1 µm.

## Discussion

The existence of a tubular axonal ER network has been known for decades. Nevertheless, the cellular mechanisms that organize a compartment that is usually distributed throughout cells, along the great lengths of axons, are until now largely unknown. The finding that several causative genes for the axon degenerative disease HSP encode ER modeling proteins, suggests a link between ER modeling and axon function or maintenance, and provides candidate proteins that may be instrumental in structure and function of the axon ER network. These candidates include several hairpin-loop-containing HSP proteins, of the spastin, atlastin, reticulon, REEP, and Arl6IP1 families, that influence ER structure in situations including yeast, mammalian cultured cells, and neuronal cell bodies in vivo (Shibata et al., 2006; Voeltz et al., 2006; Hu et al., 2008; Shibata et al., 2009; Park et al., 2010; Shibata et al., 2010). The HSP-related protein families that model ER, and some other proteins that interact with them, share a common feature of one or two intramembrane hairpin loops that can insert into the cytosolic face of the ER membrane, thereby recognizing or inducing curvature. This property makes the reticulon and REEP(DP1) families together responsible for most peripheral ER tubules in yeast, and contribute to the curved edges of ER sheets (Voeltz et al., 2006; Hu et al., 2008). The latter property may explain the expansion of ER sheets in *Drosophila* lacking the reticulon Rtnl1 (O’Sullivan et al., 2012) and in *REEP1* homozygous mutant mice (Beetz et al., 2013).

Given this background, we set out to test how far the reticulon and REEP families contribute to axonal ER organization. *REEP1* homozygous mutant mice were not previously tested for effects on axonal ER, although knockdown of *Drosophila Rtnl1* led to partial loss of smooth ER marker in distal motor axons (O’Sullivan et al., 2012). Here we build on this work by analyzing mutants of all the widely expressed and highest conserved members of the reticulon and REEP families in *Drosophila*: Rtnl1, an ortholog of all 4 human reticulons (O’Sullivan et al., 2012); ReepA, an ortholog of human REEP1-REEP4; and REEPB, an ortholog of human REEP5-REEP6. We monitored phenotypes of these mutants, singly and in combination, by visualizing ER in individual axons in situ: confocal microscopy of axonal ER markers in small numbers of motor neurons, and EM using membrane-specific staining.

Confocal microscopy revealed a partial loss of ER marker in distal but not in more proximal motor axons in *Rtnl1* and in *ReepB* mutants. Although the only *REEP* genes identified as causative for HSP are *REEP1* and *REEP2*, loss of their ortholog *ReepA* had no effect on axonal ER on its own. However, some role for ReepA in modeling axonal ER is suggested by a *ReepA*^-^ *ReepB*^-^ double mutant having a more severe axonal ER phenotype than a *ReepB*^-^ mutant (Fig. 4B). The less severe phenotype of *ReepA*^-^ compared to *ReepB*^-^ mutants is consistent with the lower levels of *ReepA* expression, judged by transcriptomics (www.flyatlas.com), and the relative expression of their GFP fusions (Fig. 2).

Given the joint and partly redundant requirement of the reticulon and REEP families for ER tubule formation in yeast (Voeltz et al., 2006), we tested whether this was also true for axonal ER. Flies lacking Rtnl1, ReepA and ReepB – equivalent to mammals lacking all four reticulons and all six REEPs, and homozygous viable – indeed showed more extreme ER fragmentation (Fig. 4) than axons lacking either Rtnl1 (Fig. 3A) or ReepA and ReepB alone (Fig. 3B). However, only a fraction of mutant axons showed this effect, and even affected axons still had continuous labeling of axons with ER marker through much of their length. ER fragmentation in the middle parts of long axons might be a consequence of axon expansion during larval growth, in which the somatic and presynaptic ends of the axon are gradually pulled apart, with insufficient ER-modeling proteins to maintain the expanding tubular network throughout the axoplasm. Therefore reticulon and REEP proteins are present in axons and have roles in ER organization there – but since triple mutant axons still mostly possess ER, there must be additional proteins required too. These might be found among the increasing number of other HSP genes that encode ER proteins with possible hairpins, such as *Arl6IP1*/*SPG61*, which affects ER organization (Novarino et al., 2014; Yamamoto et al., 2014; Fowler et al., 2016), or *C19orf12*/*SPG43* (Landouré et al., 2013). The variable nature of the triple mutant fragmentation phenotype might reflect stochastic variation in the amounts of such proteins, the amount of ER present, or external factors like physical stresses during larval movement.

EM examination of wild-type axonal ultrastructure revealed that ER tubules were effectively ubiquitous in *Drosophila* peripheral nerve sections, as seen previously in mammalian neurons (Tsukita and Ishikawa, 1976; Villegas et al., 2014), albeit with fewer tubules, presumably reflecting the smaller diameters of the axons examined here. Reconstruction over several µm showed a continuous network of ER tubules in most axons examined, in agreement with the ER continuity found in neurons by lipid dye labeling (Terasaki et al., 1994). However, in a few axons, we found short lengths of axon with no detectable ER (Fig. 6J-O). There could be several reasons for this: Terasaki et al. (1994) only assessed continuity in dendrites, cell body and proximal axon; some of the gaps we observe in EM could be short transient gaps in a dynamic network; larval axons with low diameters might be intrinsically more susceptible to occasional gaps in the ER network, than wider axons with more tubules; and we might occasionally miss an ER tubule due to weaker staining, or close proximity to other structures like plasma membrane. We also observed occasional small ER sheet-like structures in wild-type axons (Fig. 6H; Supplementary Movie 3). Although we do not see ribosomes on these, this could be due to lack of staining by the ROTO protocol. As discussed above, rough ER and translation are relatively sparse in axons, but low levels of rough ER are possible, and consistent with the occasional sheet structures observed here.

EM also showed phenotypes consistent with loss of ER membrane curvature in *Rtnl1*^-^ *ReepA*^-^ *ReepB*^-^ triple mutant axons (Fig. 6). Mutant axons had ER tubules of larger diameter, fewer tubules per axon cross-section, and consistent with our confocal data (Fig. 4), longer gaps in the ER network than wild-type, although most parts of most mutant axons examined still had a continuous ER network (Fig. 8). These mutant phenotypes could potentially have physiological consequences. Larger tubules could potentially store and release more calcium than thinner ones, while the reduced network could make the role of the ER in calcium buffering or release more localized. The less extensive ER network in mutants might also reduce the amount of contact between ER and other organelles, with consequences for calcium and lipid homeostasis that require these contacts, or for regulation of mitochondrial fission (Friedman et al., 2011) – although the continuing proximity of ER to mitochondria and plasma membrane in mutants means that any effects are presumably quantitative rather than qualitative. The reduced curvature of ER membrane in mutants might also influence their protein composition, since many membrane proteins have mechanisms for recognizing differential membrane curvature (Antonny, 2011). The occasional lack of continuity could prevent propagation of ER-dependent Ca^2+^ signals like those seen in injured mammalian sensory neurons (Cho et al., 2013); it could also cause local impairments in Ca^2+^ or lipid homeostasis that could lead to local transport inhibition, as is the case for mitochondrial transport (Wang and Schwarz, 2009). Sporadic lack of ER continuity might explain the preferential sensitivity of distal longer axons to HSPs, since these would be more likely to suffer from a gap in ER continuity to the cell body, compared to proximal or shorter axons. In this model, disease-causing alleles in single hairpin-encoding genes could promote degeneration in distal motor axons by increasing the probability of such gaps, dependent on factors such as age, axon length or diameter, and ER tubule density and dynamics.

The apparent ubiquity of ER in axons, the extent of its continuity over long distances, and the preferential susceptibility of distal longer axons to mutations that affect ER-modeling proteins, all point to important physiological roles of this compartment and of its continuity. In this work we have begun to reveal the mechanisms that determine its organization. We have shown roles for two protein families that contain HSP disease gene products, in influencing the shape of individual tubules and the axonal ER network, with potential physiological consequences that would also be affected by mutations in these genes. Understanding these physiological consequences, both in the genotypes we have described here, and in new genotypes that might also affect axonal ER organization and continuity, will provide models for the potential physiological defects in HSP and other axon degeneration diseases.

## Materials and Methods

***Drosophila* genetics.** *ReepA*^*541*^ (referred to as *ReepA*^-^), and *ReepB*^*48*^ (referred as *ReepB*^-^) mutants were generated by imprecise excision of P elements shown in Fig. 1. One of the precise excisions generated in these experiments, *ReepA*^*+C591*^ (referred to as *ReepA*^+^) was used as a genetic background control where feasible. *Rtnl1*^*1*^, referred as *Rtnl1*^-^, was a gift from G. Tear (Wakefield and Tear, 2006). *ReepA*^-^ *ReepB*^-^ and *Rtnl1*^-^ *ReepA*^-^ *ReepB*^-^ were generated by meiotic recombination on the second chromosome, and recombinants were screened for using PCR primers (Supplementary Table 2) to diagnose wild-type or mutant alleles of all three genes. Mutant and wild-type stocks were frequently genotyped to ensure that experimental flies were not contaminated. For Rtnl1 knockdown experiments, either *UAS-Rtnl1-RNAi* line 7866 (construct GD900, which has no predicted off-targets) or the *w*^*1118*^ control stock, 60000 (both obtained from the Vienna *Drosophila* RNAi Center, www.vdrc.at), was crossed with *UAS-Dcr2; CyO/If; m12-GAL4, UAS-Acsl::myc; UAS-Dcr2* (Dietzl et al., 2007) was also present for knockdown in *Rtnl1(RNAi) ReepA*^-^ *ReepB*^-^ larvae. Other fly stocks used were *P{UAS-Acsl.715.MycC}3* (Zhang et al., 2009), *PBac{681.P.FSVS-1}Rtnl1*^*CPTI001291*^ (Wakefield and Tear, 2006), *m12-GAL4* (Xiong et al., 2010; Bloomington stock 2702).

For rescue of *Rtnl1*^*1*^ we generated transgenic flies carrying *P[acman]* clone CH322-124P15 inserted at *attP2* on chromosome 3 (Bloomington stock 25710), referred to as *Rtnl1Pacman* (Venken et al., 2009). For C-terminal EGFP-LAP-tagging of ReepA and ReepB we used recombineering with the *P[acman]* system with minor modifications (Venken et al., 2009), utilizing the CH322-97D15 (*ReepA*) and CH322-16N11 (*ReepB*) BAC clones (BPRC; http://bacpac.chori.org). An EGFP-LAP-tagging cassette was amplified from R6Kamp-LAP(GFP) (Poser et al., 2008) using primers (Supplementary Table 2) with 50 bp homology to the corresponding Reep clone and around 20 bp of homology to the tagging cassette, and used to transform the recombineering *E. coli SW102* strain. Correct clones were verified at every step by PCR and/or restriction digestion. DNA for *Drosophila* transformation was extracted using a PureLink HiPure Maxiprep kit (Invitrogen), and injected into *y w M(eGFP, vasa-integrase,dmRFP)ZH-2A; M(attP)ZH-51D* or *y w M(eGFP,vasa-integrase,dmRFP)ZH-2A; M(attP) ZH-86Fb* (Stock number 24483 and 24749, Bloomington *Drosophila* Stock Center) at the Department of Genetics embryo injection facility, University of Cambridge, UK. Transformant lines were screened for the presence of the insert by PCR and GFP fluorescence. The primers used are described in Supplementary Table 2.

BLAST sequence searches were used to define genome coordinates of *P*-element excisions, and Pfam domain coordinates in coding regions, and compare protein divergence rates. They were performed at the National Center for Biotechnology Information (www.ncbi.nlm.nih.gov). REEP dendrograms were drawn from a ClustalW alignment (Larkin et al., 2007) using the neighbor-joining algorithm in MEGA 5.05 (Tamura et al., 2011).

**Histology and immunomicroscopy.** Third instar larvae were dissected in chilled Ca^2+^-free HL3 solution (Stewart et al., 1994), and fixed for 30 minutes in PBS with 4% formaldehyde. Dissected *Drosophila* preparations were permeabilized in PBS containing 0.3% Triton X-100 at room temperature, and blocked in PBT with 4% bovine serum albumin for 30 minutes at room temperature. Primary antibodies were: Csp (6D6, 1:50; Zinsmaier et al., 1994), Dlg (4F3, 1:100; Parnas et al., 2001), (both from the Developmental Studies Hybridoma Bank, Iowa, USA), GFP (Ab6556, 1:600 Abcam, UK), HRP (P7899, 1:300, Sigma), KDEL (Ab50601, 1:25, Abcam, UK), myc (2272, 1:25, Cell Signaling, USA). Fixed preparations were mounted in Vectashield (Vector Laboratories, USA), and images were collected using EZ-C1 acquisition software (Nikon) on a Nikon Eclipse C1si confocal microscope (Nikon Instruments, UK). Images were captured using 10x/0.30NA, or a 60x/1.4NA oil objective.

**Analysis of ER structure.** Confocal images were analyzed blind to genotype using ImageJ (imagej.nih.gov/ij/). Images of entire epidermal cells were obtained as z-projections of three consecutive sections. Using the line tool of ImageJ a 12-µm line was drawn from the nuclear envelope towards the periphery of each cell analyzed. Pixel intensity along the line was recorded in an Excel file. Local variance of intensity was calculated by dividing the rolling variance of the intensity (in 10-pixel windows), by the rolling mean intensity, all along the line.

Proximal (anterior) axons were imaged from segment A2, middle images were from the end of segment A4 and A5, distal (posterior) axons were imaged from segment A6 of third instar larvae. Mean gray intensity for single axon images were measured by drawing a 45-µm line, either along both M12-GAL4-expressing axons (where they could not be separated), or along the most strongly labeled axon (where they appeared as separate axons), and quantifying gray intensity (0-255) by ImageJ; occasional images with saturated pixels were excluded from analysis after blinding. Coefficient of variation was calculated by dividing the standard deviation of staining intensity by the mean; occasional images with faint staining throughout the axon were excluded from analysis after blinding. Gaps were defined as regions where staining intensity was less than 20 (out of 255), after background subtraction.

**Electron microscopy.** For epidermal cell EM, larvae were prepared and fixed as described by O’Sullivan et al. (2012). Transverse sections were cut on a Leica Ultracut UCT ultra-microtome at 70 nm, using a diamond knife, and contrasted with uranyl acetate and lead citrate (for epidermal cells). Sections were viewed using a Tecnai G2 electron microscope operated at 120 kV, and an AMT XR60B camera running the Deben software in the Multi-Imaging Centre, School of Biology, University of Cambridge.

For EM of peripheral nerves, we used a ROTO protocol (Tapia et a., 2012, Terasaki et al., 2013) to highlight membranes. Third instar larvae were dissected in HL3 solution and fixed in 0.05 M sodium cacodylate (pH 7.4) containing 4% formaldehyde, 2% vacuum distilled glutaraldehyde, and 0.2% CaCl_2_), at 4^o^C for 6 hours. Larvae were dissected as for confocal analysis, but leaving overlying organs such as gut and fat body attached, to reduce loss of peripheral nerves during processing. Preparations were then washed 3 times for 10 minutes each at 4^o^C using cold cacodylate buffer with 2 mM CaCl_2_. A solution of 3% potassium ferricyanide in 0.3 M cacodylate buffer with 4 mM CaCl_2_ was mixed with an equal volume of 4% aqueous osmium tetroxide; larval preparations were incubated in this solution at 4^o^C for 1-12 hours, then rinsed with deionized water at room temperature 5 times for 3 minutes each. Thiocarbohydrazide solution was prepared by adding 0.1 g thiocarbohydrazide to 10 ml deionized water, kept in a 60^o^C oven in a secondary embedding pot for 1 hour, swirled every 10 minutes to facilitate dissolution, and filtered through two 9 cm filter papers just before use. Larval preparations were incubated in thiocarbohydrazide solution for 20-30 minutes at room temperature and covered with foil to protect from light. Then they were rinsed with deionized water at room temperature 5 times for 3 minutes each, incubated in 2% osmium tetroxide for 30-60 minutes at room temperature, and rinsed with deionized water at room temperature 5 times for 3 minutes each. Preparations were incubated in 1% uranyl acetate (maleate-buffered to pH 5.5) at 4^o^C overnight and rinsed with deionized water at room temperature 5 times for 3 minutes each. Then they were incubated in lead aspartate solution (0.66 g lead nitrate dissolved in 100 ml 0.03 M aspartic acid, pH adjusted to 5.5 with 1 M KOH) at 60^o^C for 30 minutes and rinsed with deionized water at room temperature 5 times for 3 minutes. Then they were dehydrated twice with 50%, 70%, 90% and 100% ethanol, twice with dried ethanol, twice with dried acetone and twice with dry acetonitrile. Preparations were incubated in 50/50 acetonitrile/Quetol 651 overnight at room temperature, three times for 24 hours each in Quetol epoxy resin 651 (Agar Scientific, Stansted, UK) and three times for 24 hours each in Quetolepoxy resin 651 with BDMA (dimethylbenzylamine). They were then incubated at 60^o^C for at least 48 hours.

Serial 60-nm-thick transverse sections were cut in the larval abdominal region, visualized using scanning EM, and images were aligned for analysis of serial sections and reconstruction, as described by Terasaki et al. (2013).

**Axonal EM analyses.** To quantify axonal ER tubule diameter, non-axonal staining was removed manually, and ER tubule profiles were identified based on the local threshold in a single cross-section, and the presence of signals at the same position in adjacent sections. The minimum Feret diameter of each tubule was measured using ImageJ Fiji (https://fiji.sc) via the Analyze Particles command. ER numbers per axon were counted manually for all axons detected in the nerve. 3D reconstruction was carried out using the Fiji TrakEM2 plug-in. To quantify gaps in the tubule network, each axon was analyzed throughout the entire stack of sections. To allow for occasional lightly stained or blurred sections, only complete loss of ER tubules from three or more sections in an axon was defined as a gap. Continuous ER tubules were identified as the presence of signals at the same position for three or more sections. Given the varying brightness and contrast of EM sections, faint staining that coincided with a tubule signal in adjacent sections was also considered as an ER tubule. Color shading and sketches drawn in Fig. 7 were processed in Adobe Photoshop CS6. For quantification of glial ER sheet length, individual ER sheet profiles were measured using Fiji via the line tool and Measure command. Wrapped axons were defined as one-axon or two-axon fascicles which were completely wrapped by glial cells and isolated from other neighboring axons.

**Statistical analysis.** Statistical analyses were performed in GraphPad Prism 6. Data were analyzed by either two-tailed Student’s t-test or Mann-Whitney U tests (for data that were not normally distributed). Bar graphs and scatter plots show mean ± SEM; box plots show median with interquartile range, and the 5% and 95% percentiles as whiskers. Sample sizes are reported in figures. Confocal mean intensity datapoints from separate cells or axons were pooled across larvae or across experiments when ANOVA analysis indicated no differences among samples of the same genotype; when ANOVA analysis indicated differences among larvae or repeated experiments, mean intensity was averaged to yield datapoints for each larva or experiment, respectively. *P* levels are indicated as ns *P* > 0.05, \**P*_D_ <_D_ 0.05, \*\**P*_D_ <_D_ 0.01, \*\*\**P*_D_ <_D_ 0.001, or \*\*\*\**P*_D_ <_D_ 0.0001, except where indicated, when criteria could be defined more stringently.

## Acknowledgments

We thank Don Ryoo, Zhaohui Wang, Tony Hyman, the Developmental Studies Hybridoma Bank and the Bloomington, Vienna and Kyoto *Drosophila* Stock Centers for antibodies, bacterial constructs and stocks. We thank J. Skepper and J. Powell for help with EM, and S. Chan of the Cambridge University Genetics Department *Drosophila* facility for embryo injections.

This work was supported by grants from the Wellcome Trust (08136) and BBSRC (BB/L021706/1) to CJO’K. BY was supported by a Yousef Jameel Cambridge Trust scholarship, LZ by a Marie-Sklodowska-Curie fellowship (660516), NCO’S SZ, and OB by Marie Curie Individual Fellowships (236777, 220851 and 220874, respectively), and Z.H.K. by an A*STAR scholarship (BM/RES/07/005).

## Authorship contributions

BY, LZ and CJO’K wrote the paper. BY performed and analyzed most confocal microscopy, epidermal EM, and prepared larvae for axonal EM; LZ analyzed axonal EM data; MS generated and molecularly characterized ReepA and ReepB GFP fusions, some of the transgenic flies, and supervised generation of ReepA and ReepB mutations by ZHK, AR and MRT; NCO’S performed and analyzed epidermal EM and ER stress; BY, SZ, OB and ALP performed confocal analysis of Reep GFP fusions; MT supervised axonal EM by VB and advised on methodology and interpretation; CJO’K supervised the work.

## Supplementary Material

**Supplementary Fig. 1. Molecular lesions in *Rtnl1*^1^, *ReepA*^541^ and *ReepB*^*48*^.** BLASTN searches of the *Drosophila* genome, using the sequences of **(A)** *Rtnl1*^1^, **(B)** *ReepA*^*541*^ and **(C)** *ReepB*^*48*^ mutations as queries, show the coordinates of each molecular lesion in the genome, as gaps in alignments between mutant and genomic sequences. The *ReepA*^*541*^ and *ReepB*^*48*^ sequences highlighted in yellow show *P* element footprints left at the excision sites.

**Supplementary Fig. 2. Sketches of wrapping glia in wild-type (*ReepA*^+^) and *Rtnl1*^-^ *ReepA*^-^ *ReepB*^-^ mutant peripheral nerves, similar to those in Fig. 7J,K.** Sketches of wild-type (*ReepA*^+^, upper) and mutant (lower) wrapping glial cells were abstracted from cross-sections of peripheral nerves. Excess formation of processes were observed in triple mutants.

**Supplementary Table 1. Relative rates of evolutionary divergence of different *Drosophila* Reep proteins, shown by BLASTP searches.** BLASTP searches of Refseq proteins, using different *Drosophila melanogaster* Reep protein sequences as queries, show that CG42678 (ReepA) and CG8331 (ReepB) are diverging more slowly than CG4960, CG5539, CG11697, and CG30177, as measured by the change in E value across phylogenetic distance. Each sheet shows the hundred most similar sequences found in each BLASTP search. Hits in *D. melanogaster*, including the most closely related paralogs, are highlighted in yellow.

**Supplementary Table 2. Primers used for genotyping, *ReepA* and *ReepB* mutant screening and P[acman] C-terminal tagging.** For P[acman] tagging primers, upper case represents sequence hybridising to the gene of interest while lower case represents tagging sequence.

**Supplementary Movie 1. Serial EM sections of a wild-type peripheral nerve, showing continuity of tubular membrane structures through multiple sections.** The sixth section in the series is shown in Fig. 6A (left). ER tubules were identified as darkly stained structures present for multiple sections. See Fig. 6A for annotations.

**Supplementary Movie 2. Interactive 3D reconstruction of axonal ER from a 4.5 µm segment of a wild-type axon shown in Fig. 6F (left).** Reconstruction is generated from 75 serial 60-nm sections, and shows ER (green), mitochondria (magenta) and plasma membrane (gray).

**Supplementary Movie 3. Serial EM sections of a sheet-like ER structure in a wild-type axon, shown in Fig. 6H.** Arrows indicate short ER sheets and associated cisternae, present across multiple sections. Note that these structures usually appear continuous with tubules in adjacent sections.

**Supplementary Movie 4. Serial EM sections of a *Rtnl1*^-^ *ReepA*^-^ *ReepB*^-^ peripheral nerve showing continuity of tubular membrane structures through multiple sections.** The sixth section in the series is shown in Fig. 6A (right). ER tubules were identified as darkly stained structures present for multiple sections. See Fig. 6A for annotations.

**Supplementary Movie 5. Interactive three-dimensional reconstruction of axonal ER from a *Rtnl1*^-^ *ReepA*^-^ *ReepB*^-^ mutant axon shown in Fig. 6F (right).** Reconstruction is generated from 75 serial 60-nm sections, and shows ER (green), mitochondria (magenta) and plasma membrane (gray).

**Supplementary Movie 6. Serial EM sections of a wild-type axonal swelling shown in Fig. 6I**. An axon with a swelling is highlighted in the first frame. Sections show accumulated mitochondria, vesicles, and a large clear cisterna, as labeled in Fig. 6I.

**Supplementary Movie 7. Serial EM sections of a 4.5-µm segment of a *Rtnl1*^-^ *ReepA*^-^ *ReepB*^-^ mutant axon with disrupted continuity of ER, used for 3D reconstruction in Fig. 6K.** An axon with gaps in its ER network is highlighted in the first frame. Continuous ER tubules were identified as the presence of signals at the same position for three or more sections. Given the varying brightness and contrast of EM sections, faint staining that coincided with a tubule signal in adjacent sections was also considered as an ER tubule. Complete loss of ER tubules from three or more sections was defined as a gap.

**Supplementary Movie 8. Interactive 3D reconstruction of a 4.5-µm segment of a *Rtnl1*^-^ *ReepA*^-^ *ReepB*^-^ mutant axon with disrupted continuity of ER, shown in Fig. 6K.** The reconstruction was generated from 75 serial sections of 60 nm each. ER is in green and the plasma membrane in gray.

## References

Antonny, B. (2011). Mechanisms of membrane curvature sensing. Annu. Rev. Biochem. 80, 101–123.

Beetz, C., Pieber, T. R., Hertel, N., Schabhuttl, M., Fischer, C., Trajanoski, S., Graf, E., Keiner, S., Kurth, I., Wieland, T. et al. (2012). Exome Sequencing Identifies a REEP1 Mutation Involved in Distal Hereditary Motor Neuropathy Type V. Am. J. Hum Genet. 91, 139–145.

Beetz, C., Koch, N., Khundadze, M., Zimmer, G., Nietzsche, S., Hertel, N., Huebner, A.-K., Mumtaz, R., Schweizer, M., Dirren, E., Karle, K. N., Irintchev, A., Alvarez, V., Redies, C., Westermann, M., Kurth, I., Deufel, T., Kessels, M. M., Qualmann, B. and Hübner, C. A. (2013). A spastic paraplegia mouse model reveals REEP1-dependent ER shaping. J. Clin. Invest. 123, 4273–4282.

Ben-Yaakov, K., Dagan, S. Y., Segal-Ruder, Y., Shalem, O., Vuppalanchi, D., Willis, D. E., Yudin, D., Rishal, I., Rother, F., Bader, M., Blesch, A., Pilpel, Y., Twiss, J. L. and Fainzilber, M. (2012). Axonal transcription factors signal retrogradely in lesioned peripheral nerve. EMBO J. 31, 1350–1363.

Berridge, M. J. (1998). Neuronal calcium signaling. Neuron 21, 13–26.

Blackstone, C. (2012). Cellular pathways of hereditary spastic paraplegia. Annu. Rev. Neurosci. 35, 25–47.

Blackstone, C., O'Kane, C. J. and Reid, E. (2011). Hereditary spastic paraplegias: membrane traffic and the motor pathway. Nat. Revs. Neurosci. 12, 31–42.

Chintapalli, V. R., Wang, J. and Dow, J. A. (2007). Using FlyAtlas to identify better *Drosophila melanogaster* models of human disease. Nat. Genet. 39, 715–720.

Dietzl, G., Chen, D., Schnorrer, F., Su, K. C., Barinova, Y., Fellner, M., Gasser, B., Kinsey, K., Oppel, S., Scheiblauer, S., Couto, A., Marra, V., Keleman, K. and Dickson, B. J. (2007). A genome-wide transgenic RNAi library for conditional gene inactivation in Drosophila. Nature 448, 151–156.

Esteves T., Dürr, A., Mundwiller, M., Loureiro, J. L., Boutry, M., Gonzalez, M. A., Gauthier, J., El-Hachimi, K. H., Depienne, C., Muriel, M. P., Acosta Lebrigio, R. F., Guassen, M., Noreau, A., Speziani, F., Dionne-Laporte, A., Deleuze, J. F., Dion, P., Coutinho, P., Rouleau, G. A., Zuchner, S., Brice, A., Stevanin, G. and Darios, F. (2014). Loss of association of REEP2 with membranes leads to hereditary spastic paraplegia. Am. J. Hum. Genet. 94, 268–277.

Friedman, J. R., Lackner, L. L., West, M., DiBenedetto, J. R., Nunnari, J. and Voeltz, G. K. (2011). ER tubules mark sites of mitochondrial division. Science 334, 358–362.

Hazan, J., Fonknechten, N., Mavel, D., Paternotte, C., Samson, D., Artiguenave, F., Davoine, C. S., Cruaud, C., Dürr, A., Wincker, P. et al. (1999). Spastin, a new AAA protein, is altered in the most frequent form of autosomal dominant spastic paraplegia. Nat. Genet. 23, 296–303.

Hu, J., Shibata, Y., Voss, C., Shemesh, T., Li, Z., Coughlin, M., Kozlov, M. M., Rapoport, T. A. and Prinz, W. A. (2008). Membrane proteins of the endoplasmic reticulum induce high-curvature tubules. Science 319, 1247–1250.

Landouré, G., Zhu, P.-P., Lourenço, C. M., Johnson, J. O., Toro, C., Bricceno, K. V., Rinaldi, C., Meilleur, K. G., Sangaré, M., Diallo, O., et al. (2013). Hereditary spastic paraplegia type 43 (SPG43) is caused by mutation in C19orf12. Hum. Mutat. 34, 1357–1360.

Larkin, M. A., Blackshields, G., Brown, N. P., Chenna, R., McGettigan, P. A., McWilliam, H., Valentin, F., Wallace, I. M., Wilm, A., Lopez, R., Thompson, J. D., Gibson, T. J. and Higgins, D. G. (2007). Clustal W and Clustal X version 2.0. Bioinformatics 23, 2947–2948.

Matzat, T., Sieglitz, F., Kottmeier, R., Babatz, F., Engelen, D., and Klämbt, C. (2015). Axonal wrapping in the *Drosophila* PNS is controlled by glia-derived neuregulin homolog Vein. Development 142, 1336–1345.

Merianda, T. T., Lin, A. C., Lam, J. S., Vuppalanchi, D., Willis, D. E., Karin, N., Holt, C. E. and Twiss, J. L. (2009). A functional equivalent of endoplasmic reticulum and Golgi in axons for secretion of locally synthesized proteins. Mol. Cell. Neurosci. 40, 128–142.

Montenegro, G., Rebelo, A. P., Connell, J., Allison, R., Babalini, C., D'Aloia, M., Montieri, P., Schule, R., Ishiura, H., Price, J. et al. (2012). Mutations in the ER-shaping protein reticulon 2 cause the axon-degenerative disorder hereditary spastic paraplegia type 12. J. Clin. Invest. 122, 538–544.

O'Sullivan, N. C., Jahn, T. R., Reid, E. and O'Kane, C. J. (2012). Reticulon-like-1, the Drosophila orthologue of the Hereditary Spastic Paraplegia gene reticulon 2, is required for organization of endoplasmic reticulum and of distal motor axons. Hum. Molec. Genet. 21, 3356–3365.

Park, S. H., Zhu, P. P., Parker, R. L. and Blackstone, C. (2010). Hereditary spastic paraplegia proteins REEP1, spastin, and atlastin-1 coordinate microtubule interactions with the tubular ER network. J. Clin. Invest. 120, 1097–1110.

Parnas, D., Haghighi, A. P., Fetter, R. D., Kim, S. W. and Goodman, C. S. (2001). Regulation of postsynaptic structure and protein localization by the Rho-type guanine nucleotide exchange factor dPix. Neuron 32, 415–424.

Perry, R. B.-T., Doron-Mandel, E., Iavnilovitch, E., Rishal, I., Dagan, S. Y., Tsoory, M., Coppola, G., McDonald, M. K., Gomes, C., Geschwind, D. H., Twiss, J. L., Yaron, A. and Fainzilber, M. (2012). Subcellular knockout of importin β1 perturbs axonal retrograde signaling. Neuron 75, 294–305.

Phillips, M. J. and Voeltz, G. K. (2016). Structure and function of ER membrane contact sites with other organelles. Nat. Rev. Mol. Cell Biol. 17, 69–82.

Poser, I., Sarov, M., Hutchins, J. R., Heriche, J. K., Toyoda, Y., Pozniakovsky, A., Weigl, D., Nitzsche, A., Hegemann, B., Bird, A. W. et al. (2008). BAC TransgeneOmics: a high-throughput method for exploration of protein function in mammals. Nat. Meths. 5, 409–415.

Röper, K. (2007). Rtnl1 is enriched in a specialized germline ER that associates with ribonucleoprotein granule components. J. Cell Sci. 120, 1081–1092.

Ross, W. N. (2012). Understanding calcium waves and sparks in central neurons. Nat. Rev. Neurosci. 13, 157–168.

Ryoo, H. D., Domingos, P. M., Kang, M. J. and Steller, H. (2007). Unfolded protein response in a *Drosophila* model for retinal degeneration. EMBO J 26, 242–252.

Shibata, Y., Voeltz, G. K. and Rapoport, T. A. (2006). Rough sheets and smooth tubules. Cell 126, 435–439.

Shibata, Y., Hu, J., Kozlov, M. M. and Rapoport, T. A. (2009). Mechanisms shaping the membranes of cellular organelles. Annu. Rev. Cell Devel. Biol. 25, 329–354.

Shibata, Y., Shemesh, T., Prinz, W. A., Palazzo, A. F., Kozlov, M. M. and Rapoport, T. A. (2010). Mechanisms determining the morphology of the peripheral ER. Cell 143, 774–788.

Shibata, Y., Voss, C., Rist, J. M., Hu, J., Rapoport, T. A., Prinz, W. A. and Voeltz, G. K. (2008). The reticulon and DP1/Yop1p proteins form immobile oligomers in the tubular endoplasmic reticulum. J. Biol. Chem. 283, 18892–18904.

Shigeoka, T., Jung, H., Jung, J., Turner-Bridge, B., Ohk, J., Lin, J. Q., Amieux, P. S. and Holt, C. E. (2016) Dynamic axonal translation in developing and mature visual circuits. Cell 166, 181–192.

Stewart, B. A., Atwood, H. L., Renger, J. J., Wang, J. and Wu, C. F. (1994). Improved stability of Drosophila larval neuromuscular preparations in haemolymph-like physiological solutions. J. Comp. Physiol. A, Sensory, Neural and Behavioral Physiology 175, 179–191.

Stork, T., Engelen, D., Krudewig, A., Silies, M., Bainton, R. J. and Klämbt, C. (2008). Organization and function of the blood-brain barrier in Drosophila. J. Neurosci. 28, 587–597.

Tamura, K., Peterson, D., Peterson, N., Stecher, G., Nei, M. and Kumar, S. (2011). MEGA5: Molecular evolutionary genetic analysis using maximum likelihood, evolutionary distance, and maximum parsimony methods. Mol. Biol. Evol. 38, 2731–2739.

Tapia, J. C., Kasthuri, N., Hayworth, K. J., Schalek, R., Lichtman, J. W., Smith, S. J., and Buchanan, J. (2012). High-contrast en bloc staining of neuronal tissue for field emission scanning electron microscopy. Nat Protoc. 7, 193–206.

Terasaki, M., Slater, N. T., Fein, A., Schmidek, A. and Reese, T. S. (1994). Continuous network of endoplasmic reticulum in cerebellar Purkinje neurons. Proc. Natl. Acad. Sci. USA 91, 7510–7514.

Terasaki, M., Shemesh, T., Kasthuri, N., Klemm, R. W., Schalek, R., Hayworth, K. J., Hand, A. R., Yankova, M., Lichtman, J. W., Rapoport, T. A. and Kozlov, M. M. (2013). Stacked endoplasmic reticulum sheets are connected by helicoidal membrane motifs. Cell 154, 285–296.

Tidhar, R. and Futerman, A. H. (2013). The complexity of sphingolipid biosynthesis in the endoplasmic reticulum. Biochim. Biophys. Acta 1833, 2511–2518.

Tsukita, S. and Ishikawa, H. (1976). Three-dimensional distribution of smooth endoplasmic reticulum in myelinated axons. J. Electron Microsc. 25, 141–149.

Vance, J. E. (2015). Phospholipid synthesis and transport in mammalian cells. Traffic 16, 1–18.

Venken, K. J., Carlson, J. W., Schulze, K. L., Pan, H., He, Y., Spokony, R., Wan, K. H., Koriabine, M., de Jong, P. J., White, K. P. et al. (2009). Versatile P[acman] BAC libraries for transgenesis studies in Drosophila melanogaster. Nat. Meths. 6, 431–434.

Villegas, R., Martinez, N. W., Lillo, J., Pihan, P., Hernandez, D., Twiss, J. L. and Court, F. A. (2014). Calcium release from intra-axonal endoplasmic reticulum leads to axon degeneration through mitochondrial dysfunction. J. Neurosci. 34, 7179–7189.

Voeltz, G. K., Prinz, W. A., Shibata, Y., Rist, J. M. and Rapoport, T. A. (2006). A class of membrane proteins shaping the tubular endoplasmic reticulum. Cell 124, 573–586.

Wakefield, S. and Tear, G. (2006). The *Drosophila* reticulon, Rtnl-1, has multiple differentially expressed isoforms that are associated with a sub-compartment of the endoplasmic reticulum. Cell. Mol. Life Sci. 63, 2027–2038.

Wang, X. and Schwarz, T. L. (2009). The mechanism of Ca^2+^-dependent regulation of kinesin-mediated mitochondrial motility. Cell 136, 163–174.

Xiong, X, Wang, X., Ewanek, R., Bhat, P. and Collins, C. A. (2010). Protein turnover of the Wallenda/DLK kinase regulates a retrograde response to axonal injury. J Cell Biol. 191, 211–223.

Yamamoto, Y., Yoshida, A., Miyazaki, N., Iwasaki, K. and Sakisaka, T. (2014). Arl6IP1 has the ability to shape the mammalian ER membrane in a reticulon-like fashion. Biochem. J. 458, 68–79.

Zhang, Y., Chen, D. and Wang, Z. (2009). Analyses of mental dysfunction-related ACSL4 in *Drosophila* reveal its requirement for Dpp/BMP production and visual wiring in the brain. Hum. Molec. Genet. 18, 3894–3905.

Zhao, J., Matthies, D. S., Botzolakis, E. J., Macdonald, R. L., Blakely, R. D. and Hedera, P. (2008). Hereditary spastic paraplegia-associated mutations in the NIPA1 gene and its *Caenorhabditis elegans* homolog trigger neural degeneration in vitro and in vivo through a gain-of-function mechanism. J. Neurosci. 28, 13938–13951.

Zhao, X., Alvarado, D., Rainier, S., Lemons, R., Hedera, P., Weber, C. H., Tukel, T., Apak, M., Heiman-Patterson, T., Ming, L. et al. (2001). Mutations in a newly identified GTPase gene cause autosomal dominant hereditary spastic paraplegia. Nat. Genet. 29, 326–331.

Zinsmaier, K. E., Eberle, K. K., Buchner, E., Walter, N. and Benzer, S. (1994). Paralysis and early death in cysteine string protein mutants of Drosophila. Science 263, 977–980.

Zivraj, K. H., Tung, Y. C. L., Piper, M., Gumy, L., Fawcett, J. W., Yeo, G. S. H. and Holt, C. E. (2010). Subcellular profiling reveals distinct and developmentally regulated repertoire of growth cone mRNAs. J. Neurosci. 30, 15464–15478.

Züchner, S., Wang, G., Tran-Viet, K. N., Nance, M. A., Gaskell, P. C., Vance, J. M., Ashley-Koch, A. E. and Pericak-Vance, M. A. (2006). Mutations in the novel mitochondrial protein REEP1 cause hereditary spastic paraplegia type 31. Am. J. Hum. Genet. 79, 365–369.

